# Efficient inference of large pangenomes with PanTA

**DOI:** 10.1101/2023.07.03.547471

**Authors:** Duc Quang Le, Tien Anh Nguyen, Son Hoang Nguyen, Tam Thi Nguyen, Canh Hao Nguyen, Huong Thanh Phung, Tho Huu Ho, Nam S. Vo, Trang Nguyen, Hoang Anh Nguyen, Minh Duc Cao

## Abstract

Pangenome analysis is an indispensable step in bacterial genomics to address the high variability of bacteria genomes. However, speed and scalability remain a challenge for pangenome inference software tools to cope with the fast-growing genomic collections. We present PanTA, a software package for constructing the pangenomes of large bacterial collections. We show that PanTA exhibits an unprecedented multiple times more efficient than the current state-of-the-arts while maintaining a similar pangenome accuracy. In addition, PanTA introduces a novel mechanism to construct the pangenome progressively where new samples are added into an existing pangenome without rebuilding the accumulated collection from scratch. In the progressive mode, PanTA is demonstrated to consume orders of magnitude less computational resource than existing solutions in managing the pangenomes of growing microbial datasets. We further show that PanTA can build the pangenome of the entire collection of >28000 *Escherichia coli* genomes from the RefSeq database on a laptop computer in 32 hours, highlighting the scalability and practicality of PanTA.The software is open source and is publicly available at https://github.com/amromics/panta under an MIT license.

## Background

Prokaryotic genomes are known for enormous intraspecific variability owing to great variation events such as horizontal gene transfers, differential gene losses and gene duplication [1]. This led to the introduction of the pangenome concept as a methodology to investigate the diversity of bacterial genomes [2]. Since its inception in 2005, pangenome analysis has been a dispensable tool in microbial genomics studies [3] and has generated novel biological insights in bacterial population structures [4, 5], genetic diversity [6], niche adaptation [7] and genome assembly [8]. Pangenome studies have also been successfully applied into inferring the evolution of lineages of pandemic causing pathogens and identifying lineage-specific genetic features [9, 10], investigating genetic signatures associated with antimicrobial resistance [11], pan-reactome analyses [12], and therapeutic development including vaccine design [13] and novel drug discovery [14, 15].

To address the need for pangenome analysis, a plethora of computational tools have been developed to construct the pangenome of a collection of prokaryotic genomes. Notable examples include PGAP [16], PanOCT [17], Roary [18], BPGA [19], panX [20], MetaPGN [21], PIRATE [22], PPanGGOLiN [23], PEPPAN [24] and Panaroo [25]. The core of pangenome construction is the clustering of gene sequences into gene families. This step is typically performed by first estimating the similarity between gene sequences by a homology search tool such as CD-HIT [26], BLASTP [27] and DIAMOND [28] followed by a clustering method such as the commonly used Markov Clustering algorithm (MCL) algorithm [29]. The clustering step is also the most computationally intensive of the pipeline. The gene families are further refined through the identification of paralogous genes using either a graph-based approach or a tree-based approach. The resulting gene clusters are then classified into core or accessory genes based on their prevalence in the collection.

Advances in high-throughput sequencing technologies have recently enabled the exponential growth of microbial genomics data in public databases and in research laboratories around the world. The Genbank database stores hundreds of thousands of genomes for common bacterial species, and the numbers are fast-growing. While these resources contain rich sources of population genomics information, pangenome analysis has not been able to scale with the volume of the data. Most existing pangenome inference methods take days and require large amounts of memory that are typically beyond the capacity of a standard computer to construct the pangenome of just a few thousand isolates. In addition, the genomic databases are growing by nature, accumulating genomes of isolates collected and sequenced at different time points. There currently exists no efficient utility to update an existing pangenome when new genomes become available. In such cases, the pangenomes of the accumulated collection have to be constructed from scratch over and over, leading to the excessive burden of computational resources.

In order to address these challenges, we have developed PanTA, an efficient and scalable pangenome construction tool to keep up with the growth of bacterial genomics data sources. With vigorous computational experiments, we show that PanTA exhibits an unprecedented multiple-fold reduction in both running time and memory usage compared with the current state of the arts on building the pangenomes of large collections. Crucially, PanTA allows performing pangenome analysis progressively where batches of new samples can be added to an existing pangenome without the need to recompute the accumulated pangenome from scratch. The progressive mode can further reduce PanTA memory usage by half without affecting running time and pangenome accuracy. We also show that, PanTA in progressive mode consumed orders of magnitude less computational resource than existing solutions to manage the pangenomes of growing microbial datasets. Finally, we demonstrate the utility and practicality of PanTA by constructing the pangenome of the entire set of high quality *Escherichia coli* genomes that have been deposited into RefSeq database to date on a laptop computer.

## Results

### Overview of the pipeline

PanTA is developed with the aim to build the pangenome of a large collection of genomes, and to add a set of new genomes to an existing pangenome without rebuilding the accumulated pangenome from scratch. The workflow of PanTA pipeline is summarized in Figure 1. PanTA takes as input a list of genome assemblies and their annotations. PanTA then extracts the protein coding regions as specified by the annotation, and translates them to protein sequences. In the process, it verifies and filters out coding regions that are incorrectly annotated or that can potentially introduce noise into the clustering step and the downstream analyses.

**Figure 1:**
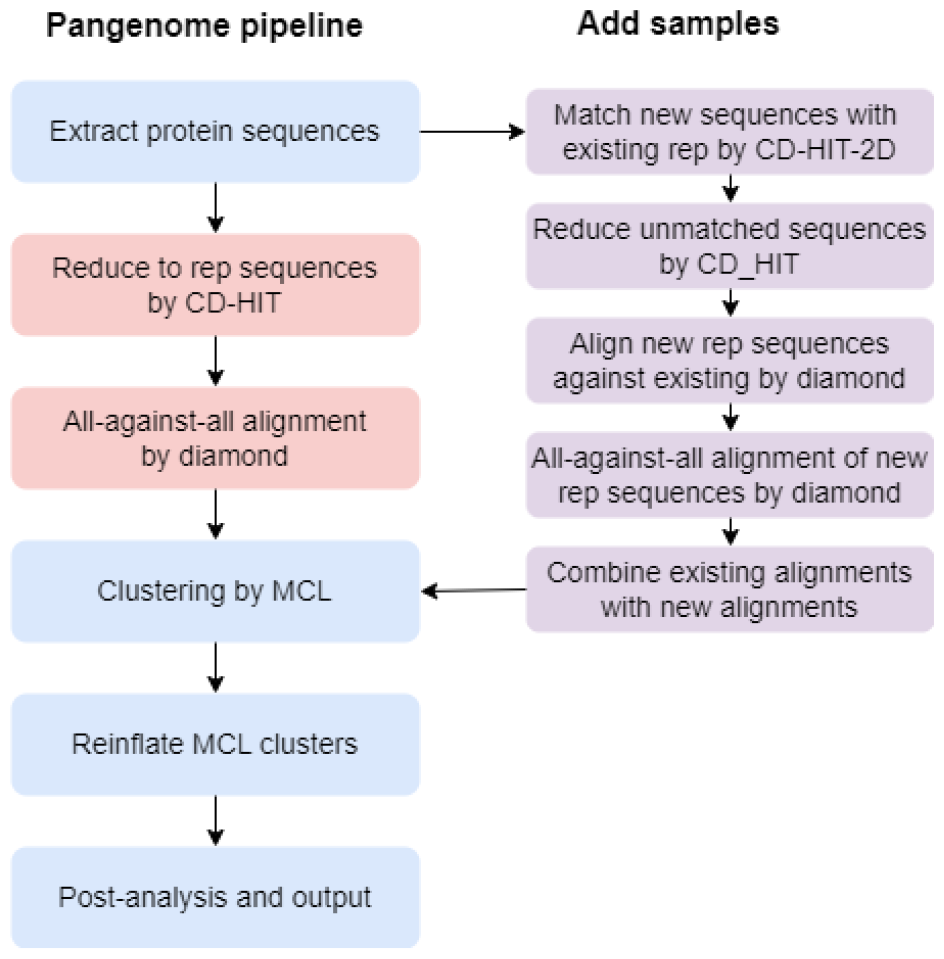
The schematic depict of PanTA workflow. The flowchart of PanTA pipeline in both single and progressive modes. In single model, the gene clustering process involves the reduction of protein sequences to representative gene sequences using CD-HIT, the all-against-all alignment of the representative sequences by DIAMOND, and the MCL clustering. In progressive mode, new protein sequences are first matched with the existing representative sequences and only unmatched sequences are reduced to form new groups. Pairwise alignments are performed only between new representative sequences against existing representative sequences and among new representative sequences.

The core of the pipeline is the clustering of all genes in the collection into gene clusters, which represent the gene families in the collection. PanTA first runs CD-HIT [26] to group similar protein sequences together, and essentially reduces the set of all protein sequences to a smaller set of representative sequences from the groups. The default identity threshold for CD-HIT grouping is 98% and the value can be adjusted by users. PanTA then performs an all-against-all alignment of the representative sequences with DIAMOND [28] or optionally BLASTP [27]. The resulting pairwise alignments are filtered to retain those that pass certain thresholds of sequence identity (default at 70%), alignment length ratios, and length difference ratios. These alignments are inputted into Markov clustering (MCL) [29] that clusters the representative sequences into homologous groups of genes. Each protein sequence is then assigned to the gene cluster its representative sequence belongs to.

While the clustering strategy employed by PanTA is similar to that of recent pangenome tools such as Roary [18], PIRATE [22] and Panaroo [25], we optimize the pipeline configurations that speed up the process without compromising the clustering accuracy. Notably, during the sequence grouping stage, PanTA runs CD-HIT only once at sequence identity 98% which is similar to Panaroo, instead of conducting multiple rounds of grouping at differing sequence identity levels as Roary and PIRATE. It also uses the word size of 5, which is suitable for such a high level of sequence identity. This word size, also used by PIRATE, enables CD-HIT to operate much faster than Panaroo’s use of word size of 2, and at the same time, produces similar sequence grouping. We also found DIAMOND significantly faster than BLASTP for the all-against-all alignment at the same level of sensitivity, confirming the previous report [28].

PanTA can run in progressive mode where it adds new genomes into an existing pangenome without rebuilding the pangenome from scratch. In this mode, PanTA uses CD-HIT-2D, a tool in the CD-HIT suite [26] to match new protein sequences extracted from the new samples to the representative sequences from the existing groups. The protein sequences that are matched to an existing group are assigned to the groups and by proxy, to the existing gene cluster. Only unmatched sequences are subject to CD-HIT to create new groups (Figure 1). By running CD-HIT clustering solely on the new genes in the added batch, PanTA significantly reduces the running time and memory usage over grouping all genes in the accumulated collection. Similarly, during the all-against-all alignment step, PanTA first performs alignment of the representative sequences of the new groups against the representative sequences of the existing groups.

It then runs the all-against-all alignment of only the new representative sequences, that is those not aligned to the existing groups to the defined sequence identity threshold. The two sets of alignments after filtering are combined and then subject to MCL clustering. With this strategy, PanTA reduces the number of sequences in the grouping and alignment steps which are the most resource-intensive steps of the whole pipeline. As a result, the process is significantly accelerated.

Finally, PanTA provides options to perform post-processing steps, including splitting paralogous clusters and multiple alignment of genes in each cluster. For split paralogs, PanTA employs the conserved gene neighborhood (CGN) approach as described in [18]. Sequences of each gene cluster are aligned using MAFFT [30] at both DNA and protein levels. PanTA then generates output reports according to the standards set out by Roary, which include a spreadsheet detailing the presence and absence of each gene in each isolate, as well as a summary of pangenome statistics.

### PanTA is significantly more efficient than existing pangenome inference tools

We evaluated the performance of PanTA and compared it with that of existing pangenome construction methods on collections of bacterial genomes. We sourced the genomes of isolates from three bacterial species *Streptococcus pneumoniae, Pseudomonas aeruginosa* and *Klebsiella pneumoniae* that are known for carrying resistance to multiple antibiotics. These three species were chosen to cover a range of genome sizes and CG content as well as both gram-positive and gram-negative. We selected 600 *S. pneumoniae*, 800 *P. aeruginosa* and 1500 *K. pneumoniae* isolates to create three datasets, named Sp600, Pa800, and Kp1500, respectively (Table 1). We downloaded their genome assemblies from the RefSeq database [31], and ran Prokka [32] to generate the gene annotations of these genomes in gff3 format. The gffs files were then used as input for the pangenome construction process.

**Table 1:** Characteristics of the three datasets to evaluate pangenome construction tools.

| Dataset | Species | Number of genomes | Genome size | Ave. gene number | CG content | Gram |
| --- | --- | --- | --- | --- | --- | --- |
| Sp600 | *S. pneumoniae* | 600 | 2.0Mb | 2.0k | 40% | positive |
| Pa800 | *P. aeruginosa* | 800 | 6.1Mb | 6.0k | 67% | negative |
| Kp1500 | *K. pneumoniae* | 1500 | 5.4Mb | 5.1k | 57% | negative |

We compared PanTA to the pangenome inference methods that are currently considered state-of-the-art in terms of scalability. Specifically, we included in the comparison Roary [18], PIRATE [22], PPanGGOLiN [23] and Panaroo [25]. Other pangenome construction methods such as panX [20], COGSoft [33] and PEPPAN [24] were reported to be prohibitively expensive for application to thousands of genomes [24, 25] and hence were excluded from the comparison. We ran all the competing tools using their default and recommended parameters. To evaluate the performance of the tools with varying input sizes, we ran them on subsets of these collections, gradually increasing in size. All computational experiments were conducted on a laptop computer with a 20 hyper-thread CPU (Intel Core i7-1280P) and 32Gb of memory, running Ubuntu Linux 22.0. All methods are parallelized with multi-threading, and we ran them on 20 threads, the number of CPU cores of the computer. We recorded the wall-time and peak memory usage of all the runs for comparison.

Most pangenome inference methods have an option to split paralogs where clusters containing paralogous genes are identified and subsequently split into true ortholog clusters. However, they have different definitions of paralogous clusters, and employ different paralog splitting strategies, leading to varying levels of splitting rigorousness. The most rigorous strategy is employed by Roary which considers a cluster paralogous if it contains more than one gene from the same genome. It then uses conserved gene neighborhood information to split homologous groups. This split paralog strategy is also performed by PanTA. PIRATE and Panaroo consider a pair of genes paralogs if they exhibit over 98% sequence identity from the CD-HIT pre-clustering step, resulting in significantly fewer paralogs compared to Roary and PanTA. PPanGGOLiN does not provide the option to split paralogs. Because of the differences in the rigorousness of the tools, we ran the competing tools with the same base configuration, that is without split paralog option. We also excluded the post-processing step that performs multiple alignment of gene clusters, as these tools eventually call a multiple alignment method such as MAFFT [30] for this task.

Figure 2a shows the computational resources in wall-time and peak memory against the size of the genome collection for the competing pangenome inference methods on the three datasets. Supplementary Figure 1 shows the differences in the number of folds in resources required by each tool against PanTA. We noted that PIRATE crashed when inferring the pangenomes for the sets of 1200 and 1500 *K. pneumoniae* genomes and PPanGGOLiN ran out of memory (32Gb) in constructing the pangenomes for 800 *P. aeruginosa* genomes and for 900 or more *K. pneumoniae* genomes. Hence, the results for these runs are not included in the comparison. We observed that all methods exhibited an approximately linear increase in time and memory usage against the input size. Strikingly, we found PanTA was significantly faster than the competing methods across three datasets by a large margin (Figure 2a, top panel). Specifically, it took under 2 minutes to build the pangenome for 600 *S. pneumoniae* genomes, and 0.168 hours and 0.207 hours to build the pangenomes for 800 *P. aeruginosa* and 1500 *K. pneumoniae* genomes, respectively. The next fastest method is PPanGGOLiN, which took between 1.8-2.2 times longer than PanTA on the small dataset Sp600, and the fold difference increased to 2.3-2.7 times in the Pa800 dataset, and 3.0-4.5 times in the Kp1500 dataset (Supplementary Figure 1). Panaroo took much longer, over 10 times longer than PanTA for the larger datasets Pa800 and Kp1500. Roary was the slowest, about 15 times slower than PanTA in most cases.

**Figure 2:**
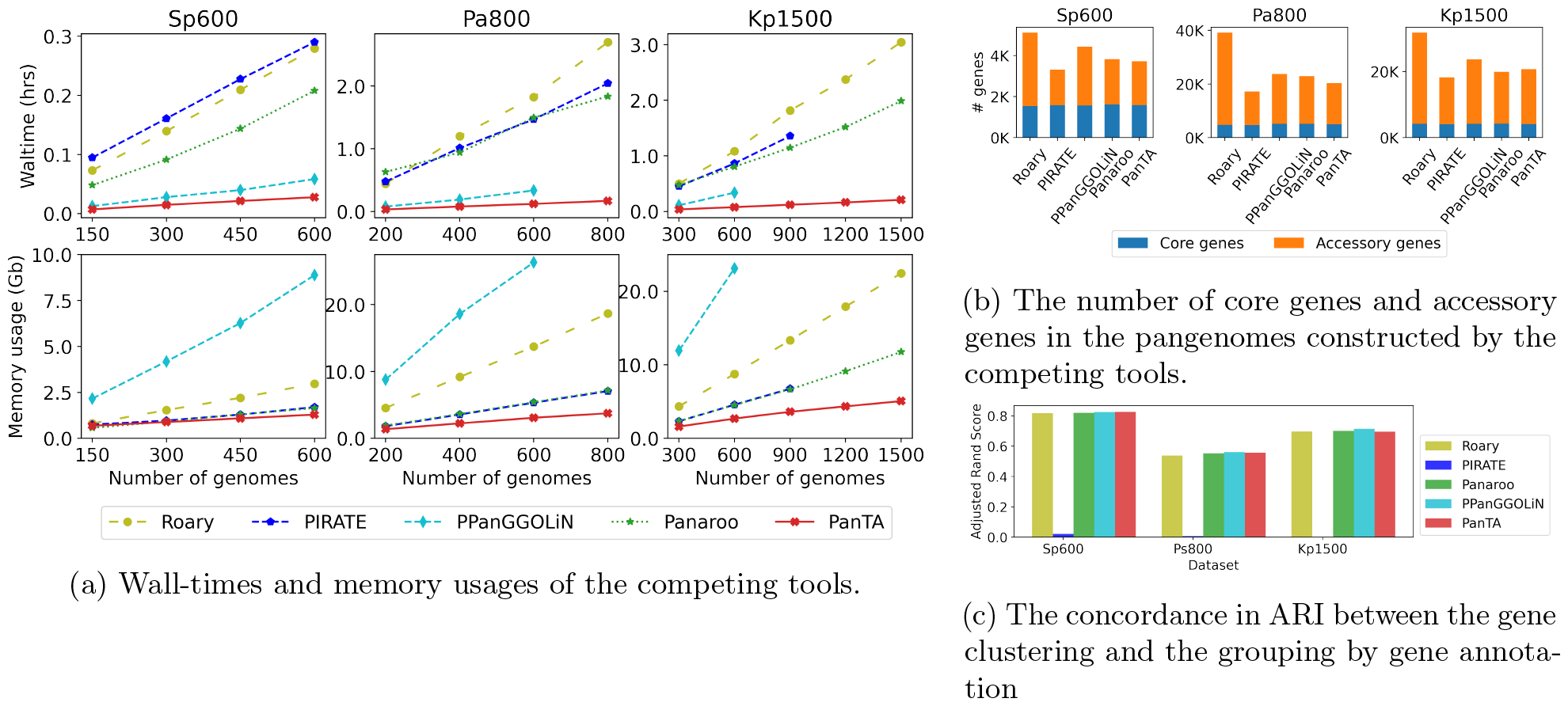
The performance of PanTA and existing tools on the three data collections. (a) Wall-time and memory usage among the competing tools at various dataset sizes. Note that PPanGGOLiN and PIRATE were unable to complete the pangenome construction for some large datasets. (b) The number of core genes and accessory genes of the pangenomes constructed by all the tools. (c) The concordance in adjusted Rand index between the gene clustering and the grouping by gene annotation.

In terms of memory usage, PanTA was also the most memory-efficient, requiring only 5.1Gb of memory for all 1500 *K. pneumoniae* genomes. Panaroo used more than twice as much memory (11.8Gb) for the same dataset, and generally the fold difference tended to increase with larger datasets. PIRATE exhibited similar memory usage profiles, but it was unable to construct the pangenomes for 1200 and 1500 *K. pneumoniae* genomes. Roary consumed 22.4Gb of memory for the Kp1500 dataset, which is 4.4 times more than PanTA. While PPanGGOLiN was the second fastest, about twice slower than PanTA, it required the most memory, about 7 times more than PanTA for the large datasets. Specifically, it required 26.3Gb and 23.1Gb of memory for analyzing 600 genomes of *P. aeruginosa* and *K. pneumoniae* respectively, it also encountered memory issues when analyzing configurations with more than 600 genomes of these species.

Figure 2b presents the numbers of core genes and of accessory genes in the pangenomes inferred by the completing methods. Note that for the Pa800 and Kp1500 datasets, PPanGGOLiN did not complete constructing the pangenomes beyond 600 genomes, we show the statistics from the pangenome constructed from the 600 genomes of each species for a fair comparison of all five methods. We observed that the pangenomes produced by Panaroo and PanTA contained a consistent number of gene families as the result of the same sequence identity threshold (70%). Roary, which used a higher threshold (95%) resulted in much higher gene clusters in its inferred pangenomes. On the other hand, PIRATE applied a series of thresholds ranging from 50% to 95% giving rise to the smallest number of gene clusters. All the methods however inferred similar numbers of core genes, in that PanTA pangenomes reported within 5% of core genes with the corresponding pangenomes produced by the other methods.

We further assessed the accuracy of the pangenomes constructed by competing methods. While there is no established benchmark to assess the accuracy of pangenome inference methods, we used the degree of concordance of the gene family clustering and the gene annotations. We collected all genes in the collection of genomes annotated by Prokka to a known gene family, that is, excluding genes that are marked as *hypothetical protein*. We considered the annotated protein names as the benchmark for evaluating the clustering of gene families. Concordance was assessed by calculating the adjusted Rand index (ARI) [34], which is a measure of similarity between clustering results. An ARI value of 1.0 indicates a perfect match between two clusterings, while a value of 0.0 indicates random grouping. The ARI of the competing methods on the three datasets is presented in Figure 2c. We found that PanTA had a similar ARI score to Panaroo, PPanGGOLiN and Roary while PIRATE performed much worse.

### PanTA progressively builds pangenome

We next evaluated the performance of PanTA in progressive mode where it updates an existing pangenome when new samples are added without the need of rebuilding the pangenome from scratch. For each of the aforementioned datasets, we ran PanTA to construct the pangenome of the smallest partition, and progressively added the genomes of the subsequent partitions into the pangenome. We noticed Panaroo also offered a similar functionality, namely Panaroo-merge, that merges the pangenomes of multiple collections together. For comparison, we ran Panaroo on each partition of the dataset, and then applied Panaroo-merge to merge the partition collections together. In these experiments, we collected the wall-times for each pangenome as the sum of the wall-time of each step, and the peak memory usage as the maximum amount of memory at each step. Figure 3 presents the computational resources consumed by both methods on the three datasets. We also included the resources needed by both methods when computing the pangenomes from scratch as part of the comparison. As presented in Figure 3, Panaroo-merge improved memory usage by 20% over the Panaroo at the cost of 70% longer running time. On the other hand, PanTA in progressive mode saved memory usage by half while maintaining a similar running time over the single mode. All in all, PanTA in progressive mode consumed only 25% and 15% of the amount of memory required by Panaroo and Panaroo-merge, respectively, while was 10 and 17 times faster.

**Figure 3:**
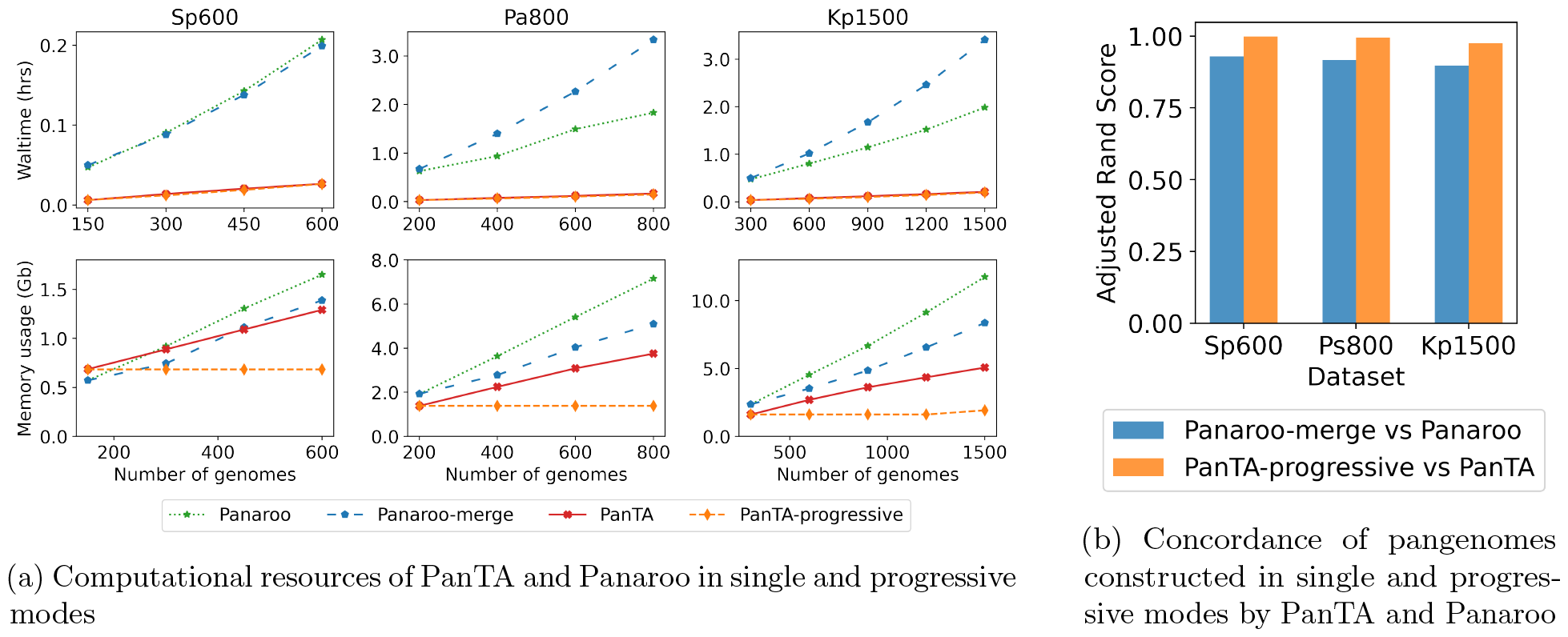
Performance of PanTA in progressive mode. (a) Comparison of computation times and memory usage of PanTA and Panaroo in single and progressive/merge modes. (b) The number of gene families and core genes inferred by PanTA and Panaroo in single and progressive modes. (c) Concordance in adjusted Rand index of pangenomes constructed in single and progressive modes by PanTA and by Panaroo. PanTA in the two modes produced near identical pangenomes while the pangenomes inferred by Panaroo and Panaroo-merge had lower concordance.

We analyzed the concordance of the pangenomes constructed by the two modes by calculating the ARIs between the two clusterings. In this calculation, we used all the genes present in the collection instead of only the annotatable genes. As shown in Figure 3b, PanTA in progressive mode produced almost identical clusterings to that in single mode (ARI*>*0.99 for Sp600 and Pa800, and *>*0.975 for the Kp1500). The pangenomes produced by the two versions of Panaroo are much less concordant, with adjusted Rand Index values of 0.93, 0.92 and 0.89 for the three datasets, respectively.

We posed a hypothetical scenario that the datasets were generated in specified batches, each at a different time point. This reflects the nature of collecting and sequencing bacterial isolates in most research laboratories, infectious disease surveillance centers and healthcare facilities. We further posited that the computational costs were measured by the time required to run on a computer with specific CPU and memory configurations, similar to those offered by a cloud computing service. We then measured the cumulative computation resources in CPU-hours required to compute the pangenomes each time a batch became available. For all methods, including Panaroo and PanTA in single mode, the computation resources would include that for recomputing the pangenomes from scratch. Panaroo-merge would only need to compute the pangenome of the new batch and then merge the pangenome of the batch to the existing pangenome. PanTA in progressive mode would add the new batch of genomes to the pangenome. Figure 4 shows the fold differences of all the methods against PanTA-progressive. As expected, PanTA-progressive required only a small fraction of computing resources compared to all other methods after a few batches. The two methods that could complete the construction of the pangenome in the Kp1500 dataset, Roary and Panaroo, respectively consumed 45.2 and 30.4 times more CPU-hours than PanTA-progressive, in addition to 11.7 and 6.1 times more memory. Although PPanGGOLiN was only 4.5 times slower than PanTA in constructing the pangenome of 600 *K. pneumoniae* genomes, the total time to compute the initial pangenome and recompute the updated pangenome was 6.9 folds higher than that of PanTA-progressive after two batches. The actual computational cost was even much higher considering that PPanGGOLiN required *>* 14.5 times as much memory. PIRATE required approximately 25-30 times more CPU-hours and 3-5 times more memory compared to PanTA-progressive after processing 3-4 batches across the three datasets. PanTA-progressive also saved 60%-70% of both CPU hours and memory uage compared to PanTA single mode.

**Figure 4:**
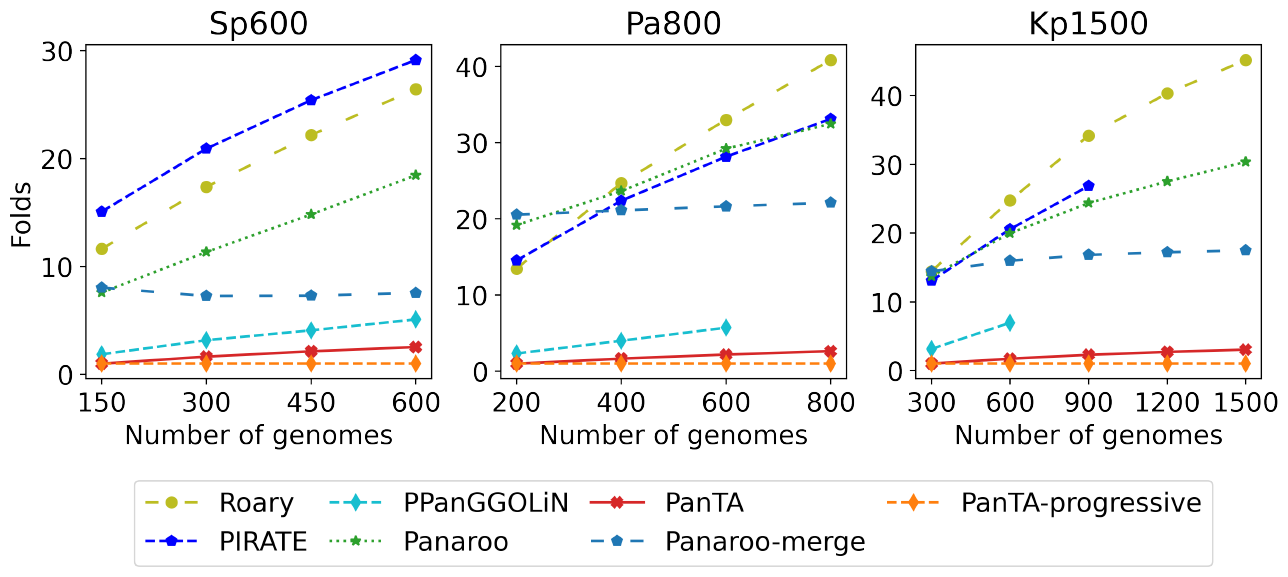
The fold difference in computational resources for both CPU-time (top panel) and memory (bottom panel) between existing methods and PanTA-progressive.

It is expected that the pangenome inference methods in single mode required higher and higher accumulated computational resources than PanTA-progressive did as more batches of data became available. We examined the resources consumed by Panaroo-merge which employs a similar approach to PanTA-progressive. Indeed, the increase of fold difference between Panaroo-merge and PanTA-progressive was much slower than other methods in single mode. However, it exhibited a large factor of fold difference, and the factor tended to increase with the genome size: 6-7X for *S. pneumoniae* (genome size 2.1Mb), 15X for *K. pneumoniae* (5.6Mb) and *>*20X for *P. aeruginosa* (6.1Mb).

### Building the pangenome of a growing genome collection

The primary goal of PanTA is to analyze and manage the extensive and fast-growing collections of microbial genomes. We demonstrate this utility by applying PanTA to a realistic and expanding collection of bacterial genomes. To this end, we collected all *Escherichia coli* genomes that were deposited into the RefSeq database [31] during the three years 2020, 2021 and 2022. *E. coli* is one of the most well-studied model prokaryotic organisms and is known for its genotypic diversity and pathogenic for both humans and animals [35]. After removing outliers, we obtained a dataset of 12,560 genomes (Methods). To demonstrate the growing nature of the dataset, we grouped the samples based on the quarters in which they were released. Table 2 shows the breakdown of the samples.

**Table 2:** Number of *E. coli* samples deposited into RefSeq database between 2020-2022 by quarter.

| Quarter | #isolates | #isolates<br>accum. |
| --- | --- | --- |
| Q1-2020 | 534 | 534 |
| Q2-2020 | 713 | 1247 |
| Q3-2020 | 830 | 2077 |
| Q4-2020 | 1109 | 3186 |
| Q1-2021 | 1166 | 4352 |
| Q2-2021 | 694 | 5046 |
| Q3-2021 | 1411 | 6457 |
| Q4-2021 | 866 | 7,323 |
| Q1-2022 | 1645 | 8,968 |
| Q2-2022 | 1233 | 10,201 |
| Q3-2022 | 1214 | 11,415 |
| Q4-2022 | 1145 | 12,560 |

We ran PanTA to build the initial pangenome of the genomes collected in the first quarter. We then progressively add genomes from subsequent quarters into the collection. For comparison with PanTA in single mode, we also ran PanTA on the accumulated data at each quarter. As shown in Figure 5, PanTA in progressive mode needed only 16.6Gb of memory to construct the pangenome for over 12000 *E. coli* genomes while the single mode consumed 30.1Gb of memory. Both modes exhibited similar running times, around 6.5 hours.

**Figure 5:**
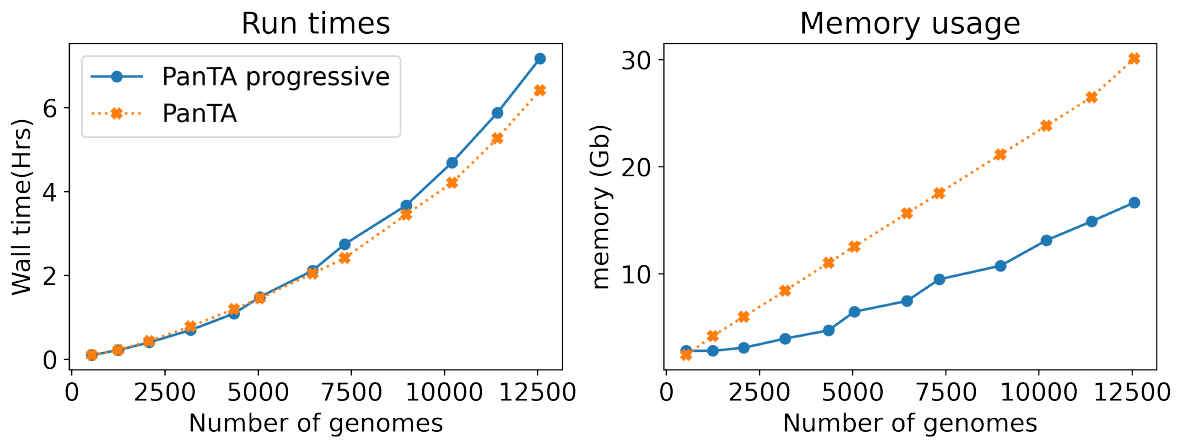
Computational resources for constructing the pangenome for *E. coli*

Encouraged by the scalability of PanTA, we proceeded to build the pangenome for the entire set of *E. coli* genomes from the RefSeq database. We downloaded all *E. coli* genomes that were released prior to 2020, and after filtering outliers, we obtained 15,625 genomes in addition to the previously collected set (Methods). We divided these genomes into batches of maximum 1000 genomes each, and iteratively added these batches into the *E. coli* pangenome with PanTA-progressive. In effect, we constructed the pangenome of all 28,275 high quality *E. coli* genomes from the RefSeq database. For this experiment, we used another laptop computer equipped with a 32-core CPU and 64Gb of memory. Strikingly, the pangenome of *E. coli* species was inferred on a laptop computer with the total time of 32 hours, including the time to build the pangenome from the past three years. The peak memory recorded during the pangenome construction was 39.9Gb.

## Discussion

Bacteria are among the most diverse life forms on earth, evidenced by the high level of variability of gene content across strains in a species. It is therefore not possible to use the genome of a single isolate as a reference genome to represent a clade. Pangenome analysis offers an alternative approach where all gene families of the clade constitute the pangenome that represents the total diversity of the clade.

Most computational methods for pangenome construction usually apply clustering of gene sequences. These methods in most cases run multiple times of CD-HIT clustering on different levels of sequence similarity in order to achieve stability of clustering. In developing PanTA, we use only one round of CD-HIT clustering and yet we obtain the near identical pangenomes with existing tools on the same sequence identity threshold. PanTA is shown to be multiple times faster than and requires less than half of the memory consumed by the current state of the arts.

The bacterial genome collections are growing by nature as more and more genomes are routinely sequenced in laboratories, as well as in research and medical settings around the world. PanTA addresses the complexity of rebuilding pangenomes by providing the progressive mode where new genomes are added into an existing pangenome. By utilizing the group membership information of the existing clustering, PanTA needs to compare the genes in the new genomes with existing groups and thereby are significantly faster than rebuilding the pangenomes. Interestingly, we found that building the pangenome progressively from batches of genomes takes a similar amount of time to build from the whole collection, and at the same time, reduces the memory requirements by half, making PanTA suitable for practical use.

The scalability of PanTA is demonstrated by the ability to construct the pangenome for a bacterial species from the entire set of *E. coli* genomes from RefSeq database on a laptop computer in an unprecedented 32-hour timeframe. PanTA can construct the pangenome progressively when new samples are added into the collection, without recomputing the accumulated collection from scratch. This further saves time and memory, and is practically suitable for analysis of the large growing collections of bacteria in the sequencing ages.

## Method

### Pangenome pipeline

PanTA accepts input genomes in GFF3 files which store gene annotations in gff format followed by the genome assembly in fasta format. This format is the output from Prokka [32] and has been popularized for the pangenome analysis started by Roary [18]. Each genome is associated with a unique ID which can be input by the user or generated by PanTA. The ID of each contig in the genome, as well as each annotated coding sequence, must be unique. Coding sequences are extracted and translated into protein sequences. Coding sequences that are less than 120 nucleotides in length or lack both a start and stop codon are excluded. Protein sequences containing more than 5% of unknown amino acids are also removed. Next, a fast sequence grouping is performed using CD-HIT [26] with an identity threshold of 98%. The representative sequences from CD-HIT are compared all-against-all by DIAMOND [28] or BLASTP [27]. The e-value threshold is set to 10e-6 by default. To reduce the time required for all-against-all alignment, the list of representative sequences is divided into smaller chunks of up to 20,000 sequences to enable parallel matching. The identified matches are filtered to retain those with sequence identity above a threshold (default at 70%). The DIAMOND result is then input into MCL [29], which uses a normalized bit score for clustering with an inflation value of 1.5. Finally, the removed sequences in the CD-HIT step are merged back into the MCL clusters. The detailed parameters of the tools are listed as follows.

- CD-HIT: cd−hit −s 0.98 −c 0.98 −T *<*number_thread*>* −M 0 −g 1 −d 256
- DIAMOND: diamond blastp −p *<*number_thread*>* −−evalue 1e−06 −−outfmt 6 qseqid sseqid pident length mismatch gapopen qstart qend sstart send evalue bitscore qlen slen −−max−target− seqs 2000
- BLASP: blastp −query *<*chunked_file*>* −db *<*blast_db*>* −evalue 10−e6 −num_threads 1 −outfmt ‘‘6 qseqid sseqid pident length mismatch gapopen qstart qend sstart send evalue bitscore qlen slen” − max_target_seqs 2000
- MCL: mcxdeblast −m9 −−score r −−line−mode=abc *<*input*>* | mcl − −−abc −I 1.5 −te *<*num_thread*>*

### Add samples pipeline

First, the protein sequences of the new samples are compared and matched with CD-HIT’s representative sequences from the previous collection. This is performed by CD-HIT-2D with the identity threshold of 98%. The protein sequences that are matched to a representative sequence are assigned to the represented group. The unmatched sequences are clustered by CD-HIT to create new groups with new representative sequences. The new representative sequences are then subject to all-against-all alignment by the alignment method of choice, *i*.*e*., DIAMOND or BLASTP. The new representative sequences are also aligned against the existing representative sequences. The two sets of alignments are then filtered according to the criteria and then combined with the existing set of alignments in the pangenome. Finally, MCL is applied to the combined set of alignments as described above.

- CD-HIT-2D cd−hit−2d −i *<*existing_group*>* −i2 *<*new_sequences*>* −s 0.98 −c 0.98 −T *<*num_threads*>* −M 0 −g 1 −d 256

### Annotating clusters

For each cluster, PanTA maintains a list of all the gene names and gene products of all genes in the cluster. It also keeps gene length statistics such as the number of genes, the minimum length, the maximum length and the average gene length in the cluster. The cluster is assigned a name taken from one of the genes that are annotated. The gene product for the cluster is the concatenation of all the gene products of the gene members. PanTA also picks the longest gene sequence to be the representative sequence for the cluster.

### Post-processing and output

PanTA presents the pangenome following the standard set out by Roary. Specifically, the presence and absence of genes in each sample are presented in a CSV format file and a Rtab format file. Upon users’ request, PanTA performs multiple sequence alignment of all gene clusters. Either or both genomic and protein sequences can be aligned. In addition, PanTA stores the existing all-against-all alignments and the existing CD-HIT groupings for subsequent analyses.

### Performance comparisons of pangenome inference methods

The lists of isolates in the three datasets Sp600, Pa800 and Kp1500, together with their Accession IDs and the URLs of their genome sequences are provided in Supporting data (see Availability of data). Their genome sequences were downloaded and were subject to annotation by Prokka in its recommended parameters (prokka −−force −−cpus 8 −−addgenes −−mincontiglen 200 −−prefix *<*accession_id *>*−−locus *<*accession_id*>* −−genus *<*genus*>* −−species *<*species*>*). The resulting annotations in gff3 format are also provided in Supporting data. The three datasets were split into batches of 150, 200 and 300 samples respectively based on the order specified in the lists.

The competing methods Roary, PIRATE, Panaroo and PPanGGOLiN were installed with their stable releases via conda. They were run with their parameters as follows:

- Roary: roary −p 20 −s −f *<*output_folder*> <*list_of_samples*>*
- PIRATE: PIRATE −−para−off −t 20 −z 0 −o *<*output folder*>* −i *<*folder_containing_samples*>*
- Panaroo: panaroo −−merge_paralogs −t 20 −−clean−mode strict −o *<*output_folder*>* −i *<*samples*>*
- Panaroo in merge mode: the pangenome for a new batch of genomes was generated with the parameters as above, and was merged into the existing pangenome with the parameters panaroo−merge −− merge_paralogs −t 20 −o *<*output_folder*>* −d *<*existing_pangenome*> <*new_batch_pangenome*>*
- PPanGGOLiN: ppanggolin workflow −−anno *<*sample list*>* −−verbose 2 −c 20 −o *<*output folder*>* −−identity 0.7
- PanTA: panta main −−dont−split −o *<*output_folder*>* −g *<*samples*>*
- PanTA in progressive mode: panta add −−dont−split −c *<*existing_pangenome*>* −g *<*samples*>*

Running times and memory usage of the computational experiments were collected with the time utility, *i*.*e*.,having /usr/bin/time −v preceding the command line. The wall time was determined from “Elapsed” field, whereas memory usage was from “Maximum resident set size”. For Panaroo in merge mode and PanTA in progressive mode, the total time of constructing the pangenome was the sum of the wall times from all preceding steps, while the memory usage was the maximum.

### Data collection for the E. coli dataset

The set of genomes available on RefSeq database was downloaded from https://ftp.ncbi.nlm.nih.gov/genomes/refseq/assembly_summary_refseq.txt (accessed Feb 22 2023). We selected only genomes of samples belonging to *E. coli* species. The genome sequence (fna file) and genome annotation (gff file) for each sample were downloaded and combined to generate a gff3 format file. Coding sequences that were shorter than 120bp or contained non-canonical nucleotides were removed. To remove outliers, we inspected the histograms of genome sizes, number of genes and N50 statistics (Supplementary Figure 2) and selected genomes that were between 4.2Mb and 5.9Mb long, contained between 4200 and 5500 genes and having N50 statistics of 50kb or higher. These genomes were grouped into quarters based on their release dates. The jupyter notebook and the script that were used to download and process the dataset and to run pangenome construction were provided in Supporting data. The resulting pangenome of the *E. coli* species was also included in Supporting data.

## Declarations

### Availability of data and materials

The PanTA software is open source and is publicly available at https://github.com/amromics/panta under an MIT license. The software is also distributed via pip. A Docker image of PanTA will be available on Docker Hub upon publishing of the manuscript. Supporting data are available on Figshare at the link https://figshare.com/s/008954ce484bb2f438f6.

## References

[1] McInerney, J.O., McNally, A., O’Connell, M.J.: Why prokaryotes have pangenomes. Nature Microbiology 2(4), 17040 (2017). doi:10.1038/nmicrobiol.2017.40

[2] Tettelin, H., Masignani, V., Cieslewicz, M.J., Donati, C., Medini, D., Ward, N.L., Angiuoli, S.V., Crabtree, J., Jones, A.L., Durkin, A.S., DeBoy, R.T., Davidsen, T.M., Mora, M., Scarselli, M., Margarit y Ros, I., Peterson, J.D., Hauser, C.R., Sundaram, J.P., Nelson, W.C., Madupu, R., Brinkac, L.M., Dodson, R.J., Rosovitz, M.J., Sullivan, S.A., Daugherty, S.C., Haft, D.H., Selengut, J., Gwinn, M.L., Zhou, L., Zafar, N., Khouri, H., Radune, D., Dimitrov, G., Watkins, K., O’Connor, K.J.B., Smith, S., Utterback, T.R., White, O., Rubens, C.E., Grandi, G., Madoff, L.C., Kasper, D.L., Telford, J.L., Wessels, M.R., Rappuoli, R., Fraser, C.M.: Genome analysis of multiple pathogenic isolates of Streptococcus agalactiae: Implications for the microbial “pan-genome”. Proceedings of the National Academy of Sciences 102(39), 13950–13955 (2005). doi:10.1073/pnas.0506758102

[3] Kim, Y., Gu, C., Kim, H.U., Lee, S.Y.: Current status of pan-genome analysis for pathogenic bacteria. Current Opinion in Biotechnology 63, 54–62 (2020). doi:10.1016/j.copbio.2019.12.001

[4] Pinto, M., González-Díaz, A., Machado, M.P., Duarte, S., Vieira, L., Carriço, J.A., Marti, S., Bajanca-Lavado, M.P., Gomes, J.P.: Insights into the population structure and pan-genome of Haemophilus influenzae. Infection, Genetics and Evolution 67, 126–135 (2019). doi:10.1016/j.meegid.2018.10.025

[5] Freschi, L., Vincent, A.T., Jeukens, J., Emond-Rheault, J.-G., Kukavica-Ibrulj, I., Dupont, M.-J., Charette, S.J., Boyle, B., Levesque, R.C.: The Pseudomonas aeruginosa Pan-Genome Provides New Insights on Its Population Structure, Horizontal Gene Transfer, and Pathogenicity. Genome Biology and Evolution 11(1), 109–120 (2019). doi:10.1093/gbe/evy259

[6] Cai, H., McLimans, C.J., Beyer, J.E., Krumholz, L.R., Hambright, K.D.: Microcystis pangenome reveals cryptic diversity within and across morphospecies. Science Advances 9(2), 1–11 (2023). doi:10.1126/sciadv.add3783

[7] Lu, Q.-F., Cao, D.-M., Su, L.-L., Li, S.-B., Ye, G.-B., Zhu, X.-Y., Wang, J.-P.: Genus-Wide Comparative Genomics Analysis of Neisseria to Identify New Genes Associated with Pathogenicity and Niche Adaptation of Neisseria Pathogens. International Journal of Genomics 2019, 1–19 (2019). doi:10.1155/2019/6015730

[8] Do, V.H., Nguyen, S.H., Le, D.Q., Nguyen, T.T., Nguyen, C.H., Ho, T.H., Vo, N.S., Nguyen, T., Nguyen, H.A., Cao, M.D.: Pasa: leveraging population pangenome graph to scaffold prokaryote genome assemblies. Nucleic Acids Research (2023). doi:10.1093/nar/gkad1170

[9] Domman, D., Quilici, M.-L., Dorman, M.J., Njamkepo, E., Mutreja, A., Mather, A.E., Delgado, G., Morales-Espinosa, R., Grimont, P.A.D., Lizárraga-Partida, M.L., Bouchier, C., Aanensen, D.M., Kuri-Morales, P., Tarr, C.L., Dougan, G., Parkhill, J., Campos, J., Cravioto, A., Weill, F.-X., Thomson, N.R.: Integrated view of Vibrio cholerae in the Americas. Science 358(6364), 789–793 (2017). doi:10.1126/science.aao2136

[10] Chung The, H., Karkey, A., Pham Thanh, D., Boinett, C.J., Cain, A.K., Ellington, M.J., Baker, K.S., Dongol, S., Thompson, C., Harris, S.R., Jombart, T., Le Thi Phuong, T., Tran Do Hoang, N., Ha Thanh, T., Shretha, S., Joshi, S., Basnyat, B., Thwaites, G., Thomson, N.R., Rabaa, M.A., Baker, S.: A high-resolution genomic analysis of multidrug-resistant hospital outbreaks of Klebsiella pneumoniae. EMBO Molecular Medicine 7(3), 227–239 (2015). doi:10.15252/emmm.201404767

[11] Kavvas, E.S., Catoiu, E., Mih, N., Yurkovich, J.T., Seif, Y., Dillon, N., Heckmann, D., Anand, A., Yang, L., Nizet, V., Monk, J.M., Palsson, B.O.: Machine learning and structural analysis of Mycobacterium tuberculosis pan-genome identifies genetic signatures of antibiotic resistance. Nature Communications 9(1), 4306 (2018). doi:10.1038/s41467-018-06634-y

[12] Seif, Y., Kavvas, E., Lachance, J.-C., Yurkovich, J.T., Nuccio, S.-P., Fang, X., Catoiu, E., Raffatellu, M., Palsson, B.O., Monk, J.M.: Genome-scale metabolic reconstructions of multiple Salmonella strains reveal serovar-specific metabolic traits. Nature Communications 9(1), 3771 (2018). doi:10.1038/s41467-018-06112-5

[13] Zeng, L., Wang, D., Hu, N., Zhu, Q., Chen, K., Dong, K., Zhang, Y., Yao, Y., Guo, X., Chang, Y.-F., Zhu, Y.: A Novel Pan-Genome Reverse Vaccinology Approach Employing a Negative-Selection Strategy for Screening Surface-Exposed Antigens against leptospirosis. Frontiers in Microbiology 8 (2017). doi:10.3389/fmicb.2017.00396

[14] Doron, S., Melamed, S., Ofir, G., Leavitt, A., Lopatina, A., Keren, M., Amitai, G., Sorek, R.: Systematic discovery of antiphage defense systems in the microbial pangenome. Science 359(6379) (2018). doi:10.1126/science.aar4120

[15] Bhardwaj, T., Somvanshi, P.: Pan-genome analysis of Clostridium botulinum reveals unique targets for drug development. Gene 623, 48–62 (2017). doi:10.1016/j.gene.2017.04.019

[16] Zhao, Y., Wu, J., Yang, J., Sun, S., Xiao, J., Yu, J.: PGAP: pan-genomes analysis pipeline. Bioinformatics 28(3), 416–418 (2012). doi:10.1093/bioinformatics/btr655

[17] Fouts, D.E., Brinkac, L., Beck, E., Inman, J., Sutton, G.: PanOCT: automated clustering of orthologs using conserved gene neighborhood for pan-genomic analysis of bacterial strains and closely related species. Nucleic Acids Research 40(22), 172–172 (2012). doi:10.1093/nar/gks757

[18] Page, A.J., Cummins, C.A., Hunt, M., Wong, V.K., Reuter, S., Holden, M.T.G., Fookes, M., Falush, D., Keane, J.A., Parkhill, J.: Roary: rapid large-scale prokaryote pan genome analysis. Bioinformatics 31(22), 3691–3693 (2015). doi:10.1093/bioinformatics/btv421

[19] Chaudhari, N.M., Gupta, V.K., Dutta, C.: BPGA-an ultra-fast pan-genome analysis pipeline. Scientific Reports 6(1), 24373 (2016). doi:10.1038/srep24373

[20] Ding, W., Baumdicker, F., Neher, R.A.: panX: pan-genome analysis and exploration. Nucleic Acids Research 46(1), 5–5 (2018). doi:10.1093/nar/gkx977

[21] Peng, Y., Tang, S., Wang, D., Zhong, H., Jia, H., Cai, X., Zhang, Z., Xiao, M., Yang, H., Wang, J., Kristiansen, K., Xu, X., Li, J.: MetaPGN: a pipeline for construction and graphical visualization of annotated pangenome networks. GigaScience 7(11), 1–11 (2018). doi:10.1093/gigascience/giy121

[22] Bayliss, S.C., Thorpe, H.A., Coyle, N.M., Sheppard, S.K., Feil, E.J.: PIRATE: A fast and scalable pangenomics toolbox for clustering diverged orthologues in bacteria. GigaScience 8(10), 1–9 (2019). doi:10.1093/gigascience/giz119

[23] Gautreau, G., Bazin, A., Gachet, M., Planel, R., Burlot, L., Dubois, M., Perrin, A., Médigue, C., Calteau, A., Cruveiller, S., Matias, C., Ambroise, C., Rocha, E.P.C., Vallenet, D.: PPanGGOLiN: Depicting microbial diversity via a partitioned pangenome graph. PLOS Computational Biology 16(3), 1007732 (2020). doi:10.1371/journal.pcbi.1007732

[24] Zhou, Z., Charlesworth, J., Achtman, M.: Accurate reconstruction of bacterial pan- and core genomes with PEPPAN. Genome Research 30(11), 1667–1679 (2020). doi:10.1101/gr.260828.120

[25] Tonkin-Hill, G., MacAlasdair, N., Ruis, C., Weimann, A., Horesh, G., Lees, J.A., Gladstone, R.A., Lo, S., Beaudoin, C., Floto, R.A., Frost, S.D.W., Corander, J., Bentley, S.D., Parkhill, J.: Producing polished prokaryotic pangenomes with the Panaroo pipeline. Genome Biology 21(1), 180 (2020). doi:10.1186/s13059-020-02090-4

[26] Li, W., Godzik, A.: Cd-hit: a fast program for clustering and comparing large sets of protein or nucleotide sequences. Bioinformatics 22(13), 1658–1659 (2006). doi:10.1093/bioinformatics/btl158

[27] Camacho, C., Coulouris, G., Avagyan, V., Ma, N., Papadopoulos, J., Bealer, K., Madden, T.L.: BLAST+: architecture and applications. BMC Bioinformatics 10(1), 421 (2009). doi:10.1186/1471-2105-10-421

[28] Buchfink, B., Xie, C., Huson, D.H.: Fast and sensitive protein alignment using DIAMOND. Nature Methods 12(1), 59–60 (2015). doi:10.1038/nmeth.3176

[29] Enright, A.J.: An efficient algorithm for large-scale detection of protein families. Nucleic Acids Research 30(7), 1575–1584 (2002). doi:10.1093/nar/30.7.1575

[30] Nakamura, T., Yamada, K.D., Tomii, K., Katoh, K.: Parallelization of MAFFT for large-scale multiple sequence alignments. Bioinformatics 34(14), 2490–2492 (2018). doi:10.1093/bioinformatics/bty121

[31] Li, W., O’Neill, K.R., Haft, D.H., DiCuccio, M., Chetvernin, V., Badretdin, A., Coulouris, G., Chitsaz, F., Derbyshire, M.K., Durkin, A.S., Gonzales, N.R., Gwadz, M., Lanczycki, C.J., Song, J.S., Thanki, N., Wang, J., Yamashita, R.A., Yang, M., Zheng, C., Marchler-Bauer, A., Thibaud-Nissen, F.: RefSeq: expanding the Prokaryotic Genome Annotation Pipeline reach with protein family model curation. Nucleic Acids Research 49(D1), 1020–1028 (2021). doi:10.1093/nar/gkaa1105

[32] Seemann, T.: Prokka: rapid prokaryotic genome annotation. Bioinformatics 30(14), 2068–2069 (2014). doi:10.1093/bioinformatics/btu153

[33] Kristensen, D.M., Kannan, L., Coleman, M.K., Wolf, Y.I., Sorokin, A., Koonin, E.V., Mushegian, A.: A low-polynomial algorithm for assembling clusters of orthologous groups from intergenomic symmetric best matches. Bioinformatics 26(12), 1481–1487 (2010). doi:10.1093/bioinformatics/btq229

[34] Rand, W.M.: Objective Criteria for the Evaluation of Clustering Methods. Journal of the American Statistical Association 66(336), 846–850 (1971). doi:10.1080/01621459.1971.10482356

[35] Tantoso, E., Eisenhaber, B., Kirsch, M., Shitov, V., Zhao, Z., Eisenhaber, F.: To kill or to be killed: pangenome analysis of Escherichia coli strains reveals a tailocin specific for pandemic ST131. BMC Biology 20(1), 146 (2022). doi:10.1186/s12915-022-01347-7

